# Orthographic deficits but typical visual perceptual processing in Chinese adults with reading disability

**DOI:** 10.1101/2023.02.13.528424

**Authors:** Xiaohui Yan, Jing Chen, Yang Fu, Yu Wu, Yixuan Ku, Fan Cao

## Abstract

Visual orthographic deficits have been reported as one of the core deficits in reading disability (RD), however, whether the deficits are orthographic-specific or domain general in all visual processing is still in debate. Hereby, we conducted an fMRI and an EEG study to examine visual orthographic deficits in Chinese adults with RD. In the fMRI study, we found that there was reduced brain activation in the left inferior temporal gyrus and right cuneus gyrus in orthographic processing (lexical minus perceptual), but not in visual perceptual processing (perceptual minus null) in adults with RD, suggesting orthographic-specific deficits. In the EEG study, adults with RD showed typical visual binding as indicated by intermodulation SSVEPs (steady-state visual-evoked potentials) for both real and pseudo characters, suggesting normal neural phase locking in the visual modality. These results consistently suggest orthographic specific deficits but normal visual perceptual processing in adults with RD, deepening our understanding of the underlying deficits associated with RD.

## Introduction

Reading disability (RD) is a specific impairment in reading which is not due to impaired cognitive ability, learning motivation or sensory acuity. It impacts about 5-10% individuals across different languages (Stevenson et al., 1982). Multiple cognitive deficits have been reported to be associated with RD in previous studies, including phonological deficits (Bruck, 1992; Hulme & Snowling, 1993), rapid automatized naming (RAN) deficits (Powell et al., 2007), auditory perceptual deficits (Richardson et al., 2004), temporal sampling deficits (Goswami, 2012), etc. Visual orthographic deficits have also been reported by many researchers (Badian, 2005; Cao et al., 2018; Cornelissen et al., 1995; Eden et al., 1996), which was even detected before reading onset (Centanni et al., 2019; Kevan & Parnmer, 2008). Orthographic intervention has also been found to promote reading (O’Brien et al., 2011). Even under the framework of the multiple deficit model, visual orthographic deficits are essential in capturing the distribution of features related to RD (Perry et al., 2019).

Although a number of studies have been conducted to understand the neurocognitive mechanisms underlying visual orthographic deficits in RD, there are many questions remained unanswered. One question concerns whether the visual orthographic deficits are linguistic-specific or domain-general. Previous studies have shown that the visual dysfunction was confined to words but not faces or other non-linguistic stimuli (Brady et al., 2021; Danelli et al., 2017). Specifically, a recent study by Brady et al. (Brady et al., 2021) found that adults with RD showed lower performance than age-controls only in a word/pseudo-word reading test but not in the Vanderbilt Holistic Face Processing Test. Another study by Danelli et al. (Danelli et al., 2017) found that adults with RD only showed deficits in a word reading task but not in non-linguistic tasks such as visual motion perception and motor learning. Functional neuroimaging studies found that orthographic deficits were associated with dysfunction of the left occipitotemporal regions (OT) and other visual regions such as the precuneus (Boros et al., 2016; Cao et al., 2018).

On the other hand, studies have reported that RD is associated with dysfunction of the magnocellular pathway (Bosse et al., 2007; Stein & Walsh, 1997). Studies have also suggested a temporal sampling deficit in RD (Goswami, 2012) which argues that the neural phase locking is reduced in individuals with RD. Evidence has been collected in the auditory modality. Specifically, the auditory cortex or other parts of the auditory pathway is entrained to the frequency of the input speech (Ding, et al., 2016), which is also called neural oscillation. However, this entrainment is reduced in individuals with DD, especially for the delta (1-3 Hz) and theta (4-8 Hz) bands (Keshavarzi, et al., 2021; Destoky, et al., 2022; Zhang, et al., 2022). However, no one has tested this hypothesis in the visual modality yet. Neural phase locking in the visual modality has been demonstrated in normal population using a paradigm of SSVEPs (steady-state visual-evoked potentials), in which stimulus in the left and right visual field flickers at two different frequencies (f1 and f2) and neural phase locking is detected at both fundamental frequencies: f1 and f2. Furthermore, neural oscillation is also detected at an intermodulation frequency, f1+f2, which is further defined as visual binding. In the current study, we examined whether individuals with RD show abnormality in neural phase locking in the visual modality using the SSVEP paradigm in order to understand whether they have low level visual deficits in addition to orthographic deficits.

The visual binding is assumed to be associated with holistic visual processing in reading. Previous studies have provided contradictory evidence in understanding whether RD is associated with deficits in holistic visual processing. One study by Conway et al. (Conway et al., 2017) found that RD readers showed no difference from control readers in a word recognition task for inverted words, but they showed a lower performance for upright words, suggesting intact visual analytic skills but impaired holistic skills in RD. In contrast, another study found that individuals with RD showed greater reliance on the whole word information when were asked to judge whether a part of the words were the same or not (Brady et al., 2021), and the result was replicated in Chinese characters (Tso et al., 2021), suggesting intact holistic processing in RD. However, the previous studies measured holistic processing indirectly and depended greatly on words/character congruency effect and observers’ subjective responses, which might cause great variance of the results. Hereby, we adopted an objective and robust measurement of holistic processing (i.e. frequency-tagged SSVEPs). Frequency-tagged SSVEPs was conducted with two parts of the visually presented object displayed at different frequencies (for example, one part flickered at f1, another at f2), and the intermodulation frequency (f1+f2) was measured as the neural signature of holistic integration. Previous study has demonstrated intermodulation frequency in typical Chinese adults using Chinese characters (Cai et al., 2020), and it was greater than that using pseudo-characters, suggesting greater holistic integration for real characters.

To answer these questions, an fMRI study with a visual spelling task and a visual symbol task and an EEG study with the frequency-tagged SSVEP task were administrated in Chinese adults with RD. We hypothesized that visual orthographic deficit is a high-ordered literacy-specific deficit rather than a low-level visual deficit, and the deficit may be related to the dysfunction of occipitoparietal regions. We also hypothesize RD readers show intact holistic processing.

### Experiment 1

#### Participants

30 University students with RD were recruited for the fMRI study. The inclusion criteria were: (1) the standard score on the Raven non-verbal IQ test was above 80, and (2) the z-score on a sentence reading fluency test or a one-minute character naming test was below -1.5. We also recruited 19 University students without RD as age-controls. The inclusion criteria were: (1) the standard score on the Raven non-verbal IQ test was above 80, and (2) the z-scores on the sentence reading fluency test and the one-minute character naming test were both above -1, and (3) the age was matched with RD. The study and consent procedures were approved by the Institutional Review Boards of the department of psychology at Sun Yat-Sen University.

After getting the informed written consent, we invited the participants for the standard Raven test as a measure of the no-verbal intelligence. A sentence reading fluency test and a one-minute character reading test were administrated as measures of reading ability. For the sentence reading fluency test, the participant was instructed to read 100 sentences and then judge whether each sentence makes sense or not within 3 minutes. For the one-minute character reading test, the participant was asked to read aloud a list of Chinese characters within one minute as fast and accurately as possible, and two lists of characters were used in the test, including a regular character list and an irregular character list. We included an initial sound deletion task, a homograph morpheme test, and a wrong character detection test to measure the phonological awareness, morphological awareness and orthographic awareness respectively. Additionally, we also included two rapid automatized naming (RAN) tests, including a digit RAN test and a picture RAN test.

#### fMRI task

A visual spelling task was employed during the MRI scanning to examine brain functional alterations during orthographic processing in Chinese adults with RD. There were three types of experimental trials in the visual spelling task, including lexical trials, non-linguistic perceptual trials, and null trails. For the lexical trials, two two-character words were visually presented sequentially, and the participant was instructed to decide whether the second characters of the pair of words shared the same radical or not. For the perceptual trials, two Tibetan symbols were visually presented sequentially, and the participant was instructed to decide whether the pair of symbols were the same or not. For the null trials, two black crosses were visually presented sequentially, and the participant was asked to press a key after the second cross onset. For each trial, the duration of each stimulus was 800 ms with a 200 ms blank between the two stimuli, as well as a 2200 to 3400 ms jittered inter-stimulus interval (ISI) following each trial. There were 96 lexical trials, 24 perceptual trials, and 48 null trials in total which were grouped into two runs. The presentation order of the trials was randomized and optimized using OptiSeq (http://surfer.nmr.mgh.harvard.edu/optseq).

#### MRI data acquisition

Brain imaging data were acquired using a 20-channel 3T Prisma Siemens scanner. The T1-weighted images were acquired using a MP-RAGE sequence, and the functional data were acquired using an EPI sequence. The parameters were as follows: for the T1-weighted images, TR=2300 ms, TE=3.24 ms, TI=900 ms, flip angle=9°, matrix size=256×256, field of view=260 mm, slice thickness=1 mm, number of slices=160; for the functional images, TR=2000 ms, TE=20 ms, flip angle=80°, matrix size=128×128, field of view=220 mm, slice thickness=3 mm, number of slices=34, and the resolution of each image was 1.7×1.7×3.0 mm.

#### fMRI data processing

The functional neuroimaging data were preprocessed using SPM12 (http://fil.ion.ucl.ac.uk/spm) with a standard pipeline. Specifically, slice timing was corrected within each volume first, and then the volumes were realigned to the first image to estimate and correct the head movement during data acquisition. When the head movement of the data exceeded 3 mm or 3°, the Art toolbox (https://www.nitrc.org/projects/artifact_detect/) was used to count how many outliers there were. If the outlier time points didn’t exceed 10% of the data, they would be scrubbed; or else, the participant was removed from further analysis. The data of one participant in the RD group was corrected and included in the further analysis. After motion correction, the data were segmented and transformed from native space into MNI space, using the parameter estimated by coregistering the structure data to the MNI template. Finally, the data were smoothed with a 4 mm full width half maximum (FWHM).

To estimate the brain functional response to the visual spelling task, a GLM model was calculated with a high pass filter of 128 seconds to reduce the linear drift. The lexical minus perceptual contrast and the perceptual minus null contrast were selected for further second level analysis. All significant results were reported at voxel level p<0.005 uncorrected, and cluster level p<0.05 FWE corrected.

## Results

### Behavioral tests and in-scanner performance

Behavioral results showed that adults with RD had poorer performance than the age controls for the following tests (t(47)=8.59, p<0.001, for the one-minute character reading test; t(47)=10.43, p<0.001, for the sentence reading fluency test; t(47)=4.36, p<0.001, for the initial sound deletion test; t(47)=2.51, p=0.016, for the homograph morpheme test; t(47)=3.08, p=0.003, for the Wrong character detection test; t(47)=-3.94, p<0.001, for digit RAN test; t(47)=-3.42, p=0.001, for picture RAN test) (Table 1).

**Table 1.** Demographic information and behavioral performance

|  | AC | RD |
| --- | --- | --- |
| N | 19 | 30 |
| Age | 249.47±26.41 | 238.4±15.03 |
| IQ | 121.68±9.89 | 116±11.16 |
| One-minute character reading test | 96.58±17.06 | 57.77±14.3*** |
| Sentence reading fluency test | 1629.58±332.23 | 794.57±228.83*** |
| Initial sound deletion test | 25.42±4.03 | 15.03±9.84*** |
| Homograph morpheme test | 27.26±1.41 | 25.47±2.91* |
| Wrong character detection test | 49.87±4.12 | 45.12±6.68** |
| Digit RAN test | 23.9±3.37 | 28.97±4.93*** |
| Picture RAN test | 41.37±6.75 | 49.35±8.62*** |
| Accuracy on the lexical trials | 0.92±0.09 | 0.91±0.11 |
| Reaction time on the lexical trials | 832.35±264.60 | 881.49±288.36 |
| Accuracy on the perceptual trials | 0.93±0.08 | 0.94±0.10 |
| Reaction time on the perceptual trials | 733.48±268.58 | 717.43±244.28 |

\* indicates $p<0.05$ , \*\* indicates $p<0.01$ , \*\*\* indicates $p<0.001$ .

For the in-scanner task, we found that adults with RD didn’t show any significant differences from the controls in accuracy or reaction time on the lexical trials (t(47)=0.24, p=0.816, for accuracy; t(47)=-0.60, p=0.552, for reaction time) or on the perceptual trials (t(47)=-0.13, p=0.897, for accuracy; t(47)=0.22, p=0.830, for reaction time).

### Neuroimaging results

To detect the group difference between adults with RD and their age-controls, two-sample t-tests were conducted for the lexical minus perceptual contrast and the perceptual minus null contrast, respectively. For the lexical minus perceptual contrast, adults with RD showed lower activation than age-controls in the left inferior temporal gyrus and right cuneus gyrus (Table 2, Figure 1), but there were no regions that showed greater activation in adults with RD than in age-controls. For the perceptual minus null contrast, there were no significant group differences.

**Table 2.** Brain regions that showed lower activation in RD than in AC for the lexical minus perceptual contrast

| Brain regions | Hemisphere |  | MNI coordinate |  |  | Z |
| --- | --- | --- | --- | --- | --- | --- |
| Inferior temporal gyrus | L | 164 | -50 | -58 | -14 | 3.91 |
|  |  |  | -42 | -54 | -2 | 3.04 |
| Cuneus | R | 170 | 12 | -70 | 28 | 3.64 |
|  |  |  | 10 | -66 | 20 | 3.36 |
|  |  |  | 10 | -78 | 30 | 3.10 |

**Figure 1.**
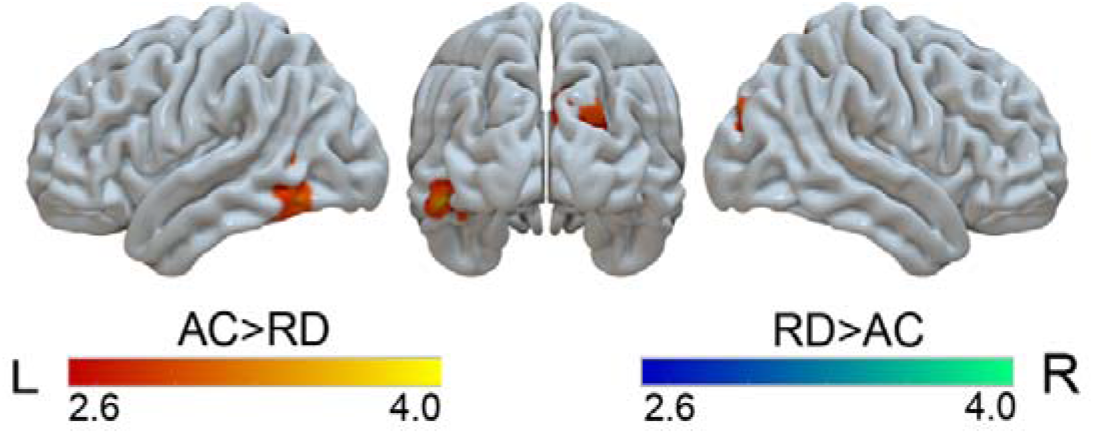
Brain regions that showed group differences in the lexical minus perceptual contrast.

### Experiment 2

#### Participants

14 University students with RD were recruited for the EEG study. The selection criteria were the same as experiment 1. Another 15 University students without RD were also recruited as age-controls.

#### SSVEPs task

A frequency-tagged SSVEP paradigm was used to examine the holistic visual processing. Specifically, 4 symmetric Chinese characters (i.e. 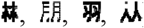) and 4 symmetric pseudo characters (i.e. (i.e. 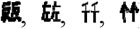)) were selected for the experiment. Each character was presented on the screen with the left half of the character flicking at 6 Hz and the right half of the character flicking at 7.5 Hz for 30s, and the on-off flicking of each half of the character was counterbalanced between left visual field and right visual field. The participant was asked to stare at a gray fixation centered on the screen, and press the space key when the fixation turned white to maintain the attention. Each trial started with a fixation, and the participant could control the presentation of the trial by pressing the space key. The trials were presented in a randomized order. The experiment was programmed with the Psychtoobox (http://psychtoolbox.org/) in Matlab.

#### EEG recoding and analysis

The EEG data were recorded continuously using a 64-channel Neuroscan system (Neuroscan, Texas, USA) with a SynAmps amplifier. The sampling rate was 1000 Hz, and electrode impedances were kept below 5 kΩ. Eye movements were monitored with two pairs of bipolar electrodes, including a pair of electrodes placed below and above the left eye to monitor the vertical movement, and a pair of electrodes placed at the left and right canthi to monitor the horizontal movement. The online reference was set at CPz.

#### EEG data processing

EEG data were processed using EEGlab and customized scripts in Matlab. Specifically, the data were off-line referenced to the average, and then segmented into epochs of 30s duration. The segmented data were detrended and multiplied by a Tukey window function (i.e., tapered cosine window, a = .02). Finally, the fast Fourier transformation (fft.m in Matlab) was conducted to transform the data into amplitude spectrum, and frequency-tagged SSVEP response was calculated. Specifically, signal-to-noise ratio (SNR) in SSVEP responses was used for further analysis. It was defined as the ratio of SSVEP response at a given frequency to its SSVEP responses at its nearby frequencies (Cai et al., 2020).

## Results

### Self-term responses

Results showed flicking characters elicited clear responses at the fundamental frequencies and several harmonics, up to the fifth harmonics (5* f1=30 Hz, 5*f2=37.5 Hz) (Figure 2). As the fifth harmonic of f1 overlapped with the forth harmonic of f2 at 30Hz, the first three harmonics were included for the further analysis. Paired t test showed that pseudo characters elicited the same self-term response as real characters (t(12) = 0.71, P = 0.491).

**Figure 2.**
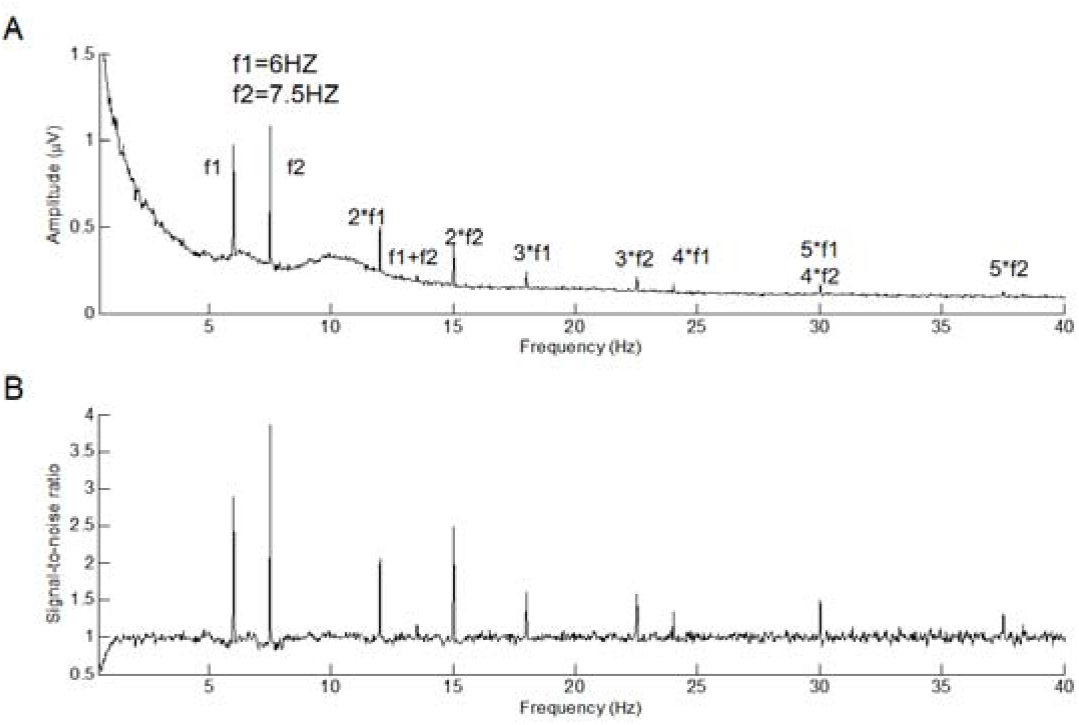
Grand-average amplitude spectrum (A) and SNR (B) of all conditions at Oz.

### Intermodulation response

For the intermodulation response, we found clear response at f1+f2 (13.5 Hz) in left occipitotemporal regions (including O1, PO7, PO5, PO3, P7, P5 and P3) (Figure 3), and these regions were selected for further analysis. Paired t test was conducted and real characters elicited greater intermodulation response than pseudo-characters (t(12) = 2.49, p = 0.027) in adults with RD (Figure 3). We also correlated the intermodulation amplitude in the real character minus pseudo character contrast with orthographic awareness, and it yielded a significant negative correlation (r=-0.54, p=0.048).

**Figure 3.**
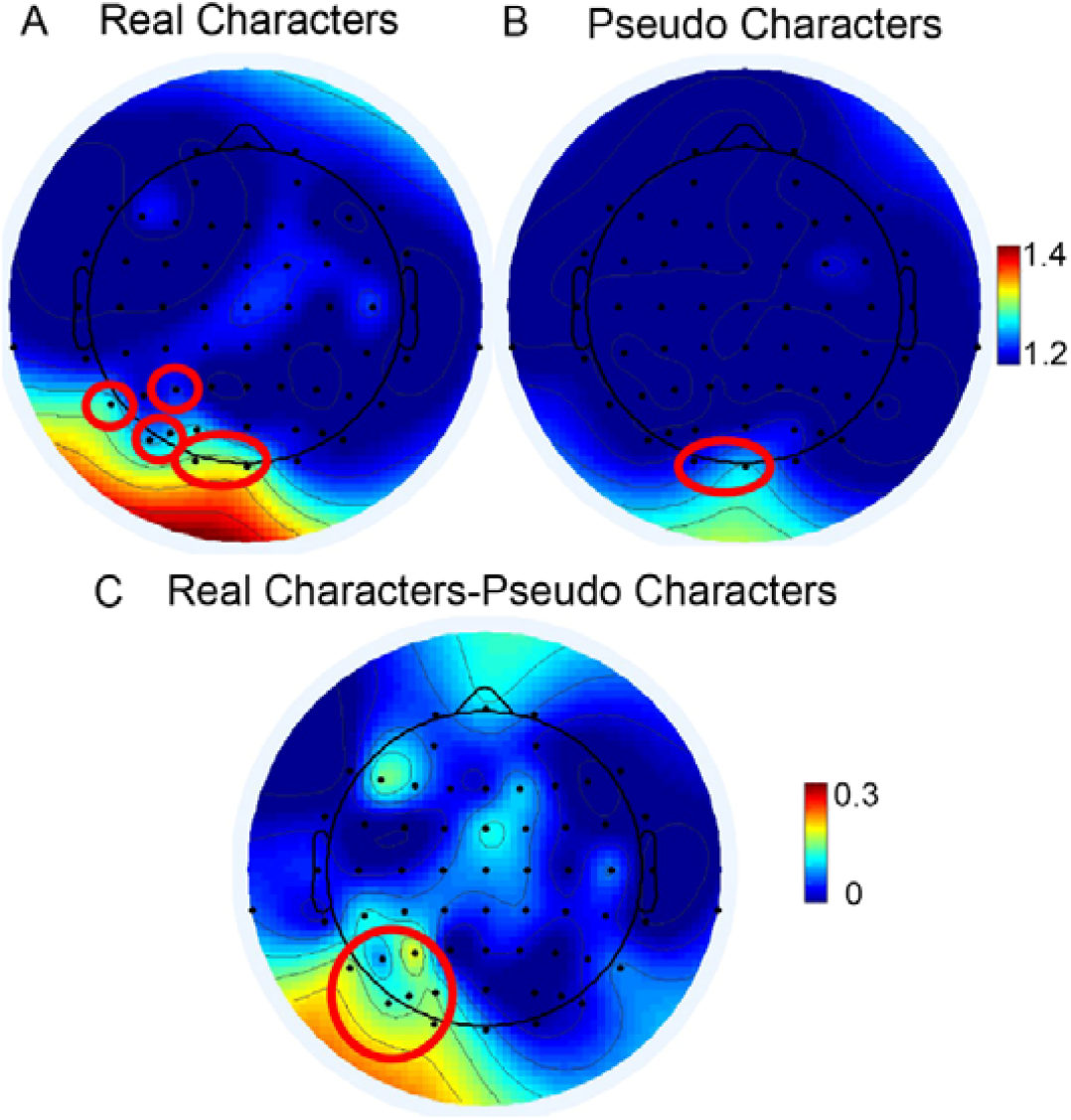
Topographic Plots for intermodulation response at f1+f2=13.5 Hz. (Read ring indicate significant response)

## Discussion

In this study, we examined the neural substrates of the visual orthographic deficits associated with adults with RD. We found that adults with RD showed lower activation in the left inferior temporal gyrus and left cuneus gyrus only in the lexical minus perceptual contrast but not in the perceptual minus null contrast, suggesting specific deficits in the orthographic processing but not in the non-linguistic visual processing. Further SSVEPs study revealed normal intermodulation responses in adults with RD. Taken together, our results suggest intact basic visual processing in adults with RD and orthographic deficits should not be due to visual problems but due to linguistic level deficits.

### Multiple deficits associated with RD

Behavioral tests revealed multiple deficits in adults with RD. Specifically, adults with RD showed lower performance on the initial sound deletion test than their age-controls, which is consistent with the phonological deficit hypothesis. Previous studies have repeatedly documented that RD was accompanied with poor phonological awareness across different ages (Snowling et al., 2019) and languages (Goswami, 2002; Siok et al., 2004). Poorer performance was also found on the wrong character detection test in adults with RD than in age-controls, suggesting visual orthographic deficits. Additional deficits were also found in the morphological awareness test and RAN tests, which is consistent with previous researches (Adrian-Ventura et al., 2020; Liu et al., 2013; Marks et al., 2022). In summary, adults with RD show deficits in phonological, orthographic, morphological and RAN processing, suggesting multiple deficits in the language domain (Perry et al., 2019).

### Visual orthographic deficits in the brain

fMRI results showed that adults with RD only had lower activation for the lexical minus perceptual contrast than age controls in left inferior temporal gyrus and right cuneus gyrus, but not for the perceptual minus null contrast, suggesting that the visuo-orthographic deficits happen at the orthographic level rather than the visual level. The results were consistent with a previous study by Danelli et al (Danelli et al., 2017), in which multiple tasks were performed and only the high-ordered literacy-related tasks revealed consistent deficits in RD. Previous research found that low-level visual deficits occurred only when compared with age-controls but not reading-level controls and phonological-based intervention could promote low-level visual skills (Olulade et al., 2013), suggesting that low-level visual deficits are consequences of RD rather than causes of RD. As a result, low-level visual deficits were not a stable feature accompanied with RD (Olulade et al., 2013). Furthermore, dysfunction of the left OT regions has been reported across tasks (Danelli et al., 2017), languages (Paulesu et al., 2001) and ages (Richlan et al., 2011), suggesting that it is a stable deficits in RD. These OT areas were important in the reading network (Pugh et al., 2000; Richlan, 2012), with the visual word form area (VWFA) located in this region (Cohen & Dehaene, 2004), and reading acquisition is accompanied by specialization of this region (Pleisch et al., 2019). However, recent studies have suggested that the left OT region is not confined to visual orthographic processing, but also involved in phonological processing (Cohen et al., 2004; Zhao et al., 2017), and an interactive region for orthography, phonology and semantics in reading prediction (Price, 2011). Therefore, we believe that deficits in the left OT region during the visual spelling task revealed in the Chinese adults with RD in the current study might reflect deficits in orthography, phonology or semantics during visual word processing.

### Intact visual holistic processing

The frequency-tagged SSVEPs study revealed normal visual binding in adults with RD as in typical controls (Cai et al., 2020), suggesting intact visual holistic processing. We also found that Chinese characters elicited greater intermodulation response than pseudo characters in adults with RD, which is also consistent with typical controls. Actually, previous studies have shown greater holistic processing in adults with RD than in controls (Brady et al., 2021; Tso et al., 2021), and the overreliance on holistic visual processing leads to inaccurate orthographic recognition.

This is consistent with our finding that there was a negative correlation between scores on the wrong character detection test and the intermodulation response. Because detecting wrong characters relies on visual analytic processing on the details of the character, rather than holistic processing, greater holistic processing is correlated with lower performance on the test.

## Conclusion

As a whole, Chinese adults with RD was accompanied with literacy-specific visual orthographic deficit, and the deficit was characterized with dysfunction of left OT regions and visual regions. However, the visual orthographic deficit was not caused by deficient holistic processing.

## Reference

Adrian-Ventura, J., Soriano-Ferrer, M., Fuentes-Claramonte, P., Morte-Soriano, M., Parcet, M. A., & Avila, C. (2020). Grey matter reduction in the occipitotemporal cortex in Spanish children with dyslexia: A voxel-based morphometry study. Journal of Neurolinguistics, 53. doi:10.1016/j.jneuroling.2019.100873

Badian, N. A. (2005). Does a visual-orthographic deficit contribute to reading disability? Annals of Dyslexia, 55(1), 28–52. doi:10.1007/s11881-005-0003-x

Boros, M., Anton, J. L., Pech-Georgel, C., Grainger, J., Szwed, M., & Ziegler, J. C. (2016). Orthographic processing deficits in developmental dyslexia: Beyond the ventral visual stream. Neuroimage, 128, 316–327.

Bosse, M. L., Tainturier, M. J., & Valdois, S. (2007). Developmental dyslexia: the visual attention span deficit hypothesis. Cognition, 104(2), 198–230. doi:10.1016/j.cognition.2006.05.009

Brady, N., Darmody, K., Newell, F. N., & Cooney, S. M. (2021). Holistic processing of faces and words predicts reading accuracy and speed in dyslexic readers. PLoS One, 16(12), e0259986. doi:10.1371/journal.pone.0259986

Bruck, M. (1992). Persistence of dyslexics’ phonological deficits. Developmental Psychology, 28(5), 874–886.

Cai, Y., Mao, Y., Ku, Y., & Chen, J. (2020). Holistic Integration in the Processing of Chinese Characters as Revealed by Electroencephalography Frequency Tagging. Perception, 49(6), 658–671. doi:10.1177/0301006620929197

Cao, F., Yan, X., Spray, G. J., Liu, Y., & Deng, Y. (2018). Brain mechanisms underlying visuo-orthographic deficits in children with developmental dyslexia. Frontiers in Human Neuroscience, 12, 490. doi:10.3389/fnhum.2018.00490

Centanni, T. M., Norton, E. S., Ozernov-Palchik, O., Park, A., Beach, S. D., Halverson, K., Gaab, N., & Gabrieli, J. D. E. (2019). Disrupted left fusiform response to print in beginning kindergartners is associated with subsequent reading. Neuroimage Clin, 22, 101715. doi:10.1016/j.nicl.2019.101715

Cohen, L., Jobert, A., Le Bihan, D., & Dehaene, S. (2004). Distinct unimodal and multimodal regions for word processing in the left temporal cortex. Neuroimage, 23(4), 1256–1270. doi:10.1016/j.neuroimage.2004.07.052

Cohen, L., & Dehaene, S. (2004). Specialization within the ventral stream: the case for the visual word form area. Neuroimage, 22(1), 466–476. doi:10.1016/j.neuroimage.2003.12.049

Conway, A., Brady, N., & Misra, K. (2017). Holistic word processing in dyslexia. PLoS One, 12(11), e0187326. doi:10.1371/journal.pone.0187326

Cornelissen, P., Richardson, A., Mason, A., Fowler, S., & Stein, J. (1995). Contrast sensitivity and coherent motion detection measured at photopic luminance levels in dyslexics and controls. Vision Research, 35(10), 1483–1494. doi:10.1016/0042-6989(95)98728-R

Danelli, L., Berlingeri, M., Bottini, G., Borghese, N. A., Lucchese, M., Sberna, M., Price, C. J., & Paulesu, E. (2017). How many deficits in the same dyslexic brains? A behavioural and fMRI assessment of comorbidity in adult dyslexics. Cortex, 97, 125–142.

Eden, G. F., Stein, J. F., Wood, H. M., & Wood, F. B. (1996). Differences in visuospatial judgement in reading-disabled and normal children. Percept Mot Skills, 82(1), 155–177. doi:10.2466/pms.1996.82.1.155

Goswami, U. (2002). Phonology, reading development, and dyslexia: A cross-linguistic perspective. Annals of Dyslexia, 52, 141–163.

Hulme, C., & Snowling, M. (1993). Phonological deficits and the development of word recognition skills in developmental dyslexia. Reading Disabilities : Diagnosis and Component Processes, 74, 225–236.

Kevan, A., & Parnmer, K. (2008). Visual deficits in pre-readers at familial risk for dyslexia. Vision Research, 48(28), 2835–2839. doi:10.1016/j.visres.2008.09.022

Liu, L., Tao, R., Wang, W. J., You, W. P., Peng, D. L., & Booth, J. R. (2013). Chinese dyslexics show neural differences in morphological processing. Dev Cogn Neurosci, 6, 40–50. doi:10.1016/j.dcn.2013.06.004

Marks, R. A., Eggleston, R. L., Sun, X., Yu, C.-L., Zhang, K., Nickerson, N., Hu, X.-S., & Kovelman, I. (2022). The neurocognitive basis of morphological processing in typical and im paired readers. Annals of Dyslexia, 72(2), 361–383. doi:10.1007/s11881-021-00239-9

O’Brien, B. A., Wolf, M., Miller, L. T., Lovett, M. W., & Morris, R. (2011). Orthographic processing efficiency in developmental dyslexia: an investigation of age and treatment factors at the sublexical level. Annals of Dyslexia, 61(1), 111–135. doi:10.1007/s11881-010-0050-9

Olulade, O. A., Napoliello, E. M., & Eden, G. F. (2013). Abnormal visual motion processing is not a cause of dyslexia. Neuron, 79(1), 180–190. doi:10.1016/j.neuron.2013.05.002

Paulesu, E., ., Démonet, J. F., Fazio, F., ., Mccrory, E., ., Chanoine, V., ., Brunswick, N., ., Cappa, S. F., Cossu, G., ., Habib, M., ., & Frith, C. D. (2001). Dyslexia: Cultural diversity and biological unity. Science, 291(5511), 2165–2167.

Perry, C., Zorzi, M., & Ziegler, J. C. (2019). Understanding dyslexia through personalized large-scale computational models. Psychological Science, 30(3), 386–395. doi:10.1177/0956797618823540

Pleisch, G., Karipidis, II, Brauchli, C., Rothlisberger, M., Hofstetter, C., Stampfli, P., Walitza, S., & Brem, S. (2019). Emerging neural specialization of the ventral occipitotemporal cortex to characters through phonological association learning in preschool children. Neuroimage, 189, 813–831. doi:10.1016/j.neuroimage.2019.01.046

Powell, D., Stainthorp, R., Stuart, M., Garwood, H., & Quinlan, P. (2007). An experimental comparison between rival theories of rapid automatized naming performance and its relationship to reading. Journal of Experimental Child Psychology, 98(1), 46–68. doi:10.1016/j.jecp.2007.04.003

Pugh, K. R., Mencl, W. E., Jenner, A. R., Katz, L., Frost, S. J., Lee, J. R., Shaywitz, S. E., & Shaywitz, B. A. (2000). Functional neuroimaging studies of reading and reading disability (developmental dyslexia). Mental Retardation and Developmental Disabilities Research Reviews, 6(3), 207–213. doi:10.1002/1098-2779(2000)6:3<207::Aid-Mrdd8>3.3.Co;2-G

Richardson, U., Thomson, J. M., Scott, S. K., & Goswami, U. (2004). Auditory processing skills and phonological representation in dyslexic children. Dyslexia, 10(3), 215–233. doi:10.1002/dys.276

Richlan, F., Kronbichler, M., & Wimmer, H. (2011). Meta-analyzing brain dysfunctions in dyslexic children and adults. Neuroimage, 56(3), 1735–1742. doi:10.1016/j.neuroimage.2011.02.040

Richlan, F. (2012). Developmental dyslexia: dysfunction of a left hemisphere reading network. Frontiers in Human Neuroscience, 6, 120. doi:10.3389/fnhum.2012.00120

Siok, W. T., Perfetti, C. A., Jin, Z., & Tan, L. H. (2004). Biological abnormality of impaired reading is constrained by culture. Nature, 431(7004), 71–76. doi:10.1038/nature02865

Snowling, M. J., Lervag, A., Nash, H. M., & Hulme, C. (2019). Longitudinal relationships between speech perception, phonological skills and reading in children at high-risk of dyslexia. Developmental Science, 22(1). doi:10.1111/desc.12723

Stein, J., & Walsh, V. (1997). To see but not to read; the magnocellular theory of dyslexia. Trends in Neurosciences, 20(4), 147–152. doi:10.1016/s0166-2236(96)01005-3

Stevenson, H. W., Stigler, J. W., Lucker, G. W., Lee, S., Hsu, C., & Kitamura, S. (1982). Reading disabilities: the case of Chinese, Japanese, and English. Child Development, 53(5), 1164–1181.

Tso, R. V., Chan, R. T., Chan, Y. F., & Lin, D. (2021). Holistic processing of Chinese characters in college students with dyslexia. Sci Rep, 11(1), 1973. doi:10.1038/s41598-021-81553-5

Zhao, L. B., Chen, C. H., Shao, L. Y., Wang, Y. P., Xiao, X. Q., Chen, C. S., Yang, J. F., Zevin, J., & Xue, G. (2017). Orthographic and phonological representations in the fusiform cortex. Cerebral Cortex, 27(11), 5197–5210. doi:10.1093/cercor/bhw300

